# Polyserine peptides are toxic and exacerbate tau pathology in mice

**DOI:** 10.1101/2024.10.10.616100

**Authors:** Meaghan Van Alstyne, Vanessa L. Nguyen, Charles A. Hoeffer, Roy Parker

## Abstract

Polyserine domains mediate the association of nuclear RNA binding proteins with cytoplasmic tau aggregates that occurs across tauopathy models and patient samples. In cell lines, polyserine peptides co-localize with and promote formation of tau aggregates suggesting the cytoplasmic mislocalization of polyserine-containing proteins might contribute to human disease. Moreover, polyserine can be produced by repeat associated non-AUG translation in CAG repeat expansion diseases. However, whether polyserine expressed in a mammalian brain is toxic and/or can exacerbate tau pathology is unknown. Here, we used AAV9-mediated delivery to express a 42-repeat polyserine protein in wild-type and tau transgenic mouse models. We observe that polyserine expression has toxic effects in wild-type animals indicated by reduced weight, behavioral abnormalities and a striking loss of Purkinje cells. Moreover, in the presence of a pathogenic variant of human tau, polyserine exacerbates disease markers such as phosphorylated and insoluble tau levels and the seeding capacity of brain extracts. These findings demonstrate that polyserine domains can promote tau-mediated pathology in a mouse model and are consistent with the hypothesis that cytoplasmic mislocalization of polyserine containing proteins might contribute to the progression of human tauopathies.

## Introduction

The microtubule-binding protein tau is a key player in several neurodegenerative diseases termed tauopathies.^1^ In disease, tau transitions into an insoluble state, forming intracellular aggregates – a hallmark feature of pathology.^1^ Multiple factors can contribute to tau pathobiology such as post-translational modifications or disease-associated mutations that promote transition from a microtubule bound state to an aggregated fibrillar form characteristic of disease.^2^ Moreover, *in vitro* polyanionic co-factors such as RNA accelerate formation of fibrillar aggregates.^3,4^ However, how potential cellular co-factors may affect tau fibrillization in mammalian neurons remains incompletely understood.

We and others, have demonstrated that RNA binding proteins such as SRRM2 and Pinin mislocalize to tau aggregates in cell models, animal models and post-mortem patient tissue.^5,6^ Long-stretches of serine repeats or serine-rich regions drive this association with tau.^7^ Interestingly, in cellular models, polyserine (polySer)-containing proteins or exogenous polySer form assemblies that are preferred sites of tau aggregation.^7^ Further, the levels of polySer-containing proteins correlate with the extent of tau aggregation.^7^ However, whether polySer modulates tau aggregation and pathology in animal models is unknown.

In addition to being contained within proteins, polySer peptides can be endogenously expressed in the context of repeat expansion diseases. Specifically, in spinocerebellar ataxia 8 (SCA8) polySer containing proteins are expressed through repeat associated non-AUG (RAN) translation of transcripts derived from a CAG expansion translating in the AGC frame.^8^ In this context, polySer has been shown to accumulate in white matter regions where it coincides with demyelination and degeneration.^8^ Moreover, polySer proteins are also expressed from the CAG expansion casual in Huntington’s disease (HD) and are present in striatal regions prominently affected in disease.^9,10^ Thus, polySer proteins are expressed across repeat expansion diseases where they may contribute to neurotoxicity. However, the toxic properties of polySer in the mammalian brain – either dependent or independent of tau – remain incompletely understood.

Here, we expressed polySer in wild-type and tau transgenic mouse models to determine both the effects of polySer expression alone and the potential modulation of tau pathology. We observe that polySer induces behavioral abnormalities in wild-type mice that coincide with a loss of Purkinje cells. In addition, polySer exacerbates disease-relevant markers in tau transgenic mice. These observations provide evidence of polySer toxicity *in vivo* that may contribute to the pathogenesis of specific repeat expansion diseases. Further, they demonstrate that levels of polySer modulate tau pathology in an animal model, consistent with the hypothesis that increased associations of polySer-containing proteins and tau could contribute to the progression of human tauopathies.

## Results

### AAV9-mediated expression of polyserine induces toxicity in wild-type and tau transgenic mice

To investigate the effect of polySer *in vivo*, we utilized AAV9 gene delivery to express a GFP control or a GFP-tagged repeat of 42 serines (Ser_42_), the length of the longest polySer stretch in SRRM2.^5,7^ AAV9 was delivered to the central nervous system (CNS) by intracerebroventricular injection at P1 to both wild-type (WT) and tau mice harboring a 1N4R human tau transgene with a P301S disease-linked mutation (PS19) (Figure 1A).^11^

**Figure 1.**
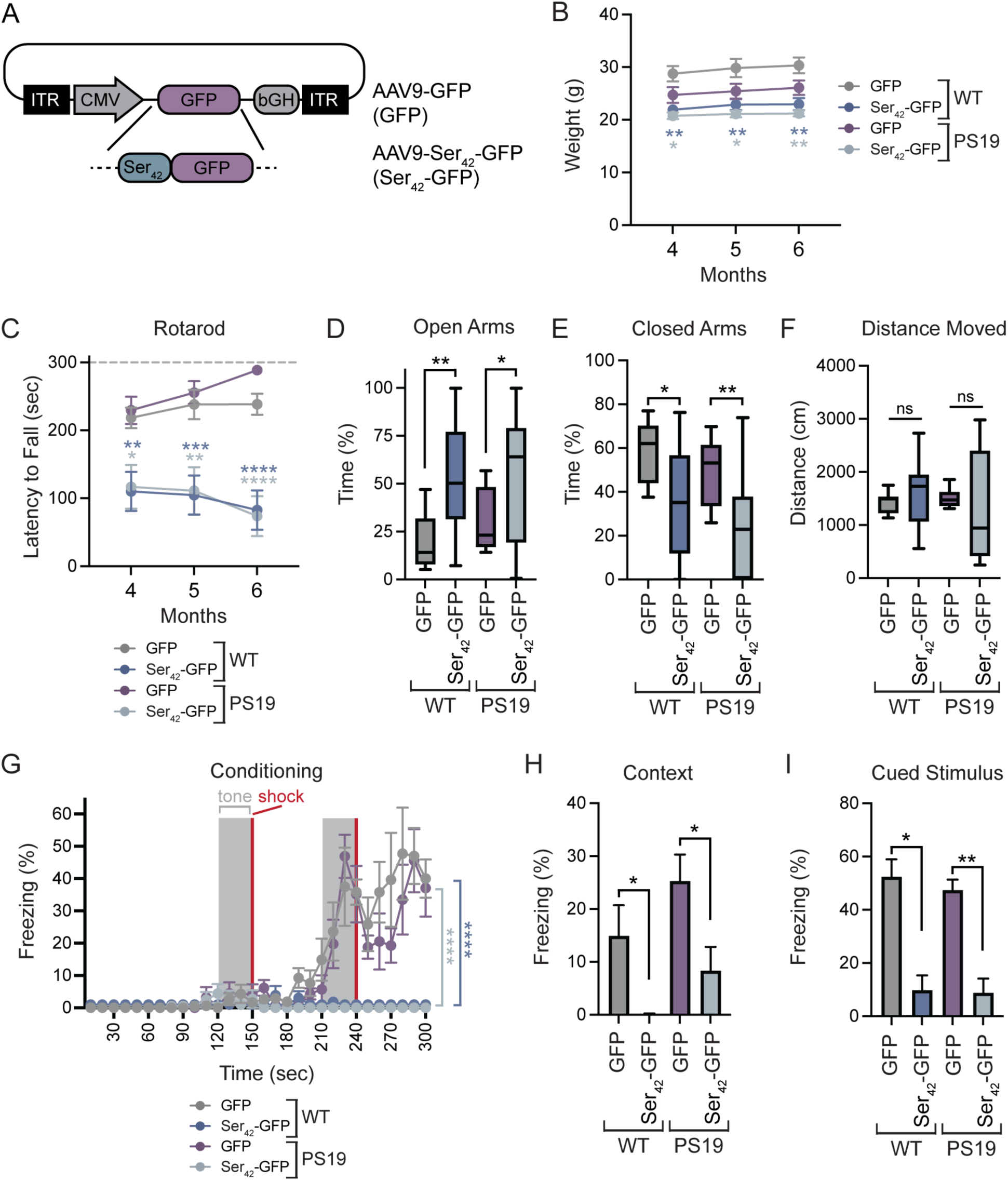
AAV9-mediated overexpression of polyserine leads to motor dysfunction in wild-type and tau transgenic mice. (**A**) Schematic of AAV constructs for *in vivo* expression of transgenes. (**B**) Weight of WT and PS19 tau transgenic animals injected with 1×10^11^ vgs of AAV9-GFP or AAV9-Ser_42_-GFP per animal by ICV injection at P1. Data represents mean and SEM. n>=8 animals per group. Statistics performed between GFP and Ser_42_-GFP injected animals within paired genotypes with two-way ANOVA and Bonferroni’s correction. (**C**) Latency to fall on rotarod assay of WT and PS19 animals injected with AAV9 as in (B). Data represents mean and SEM. n>=12 animals per group. Statistics performed between GFP and Ser_42_-GFP injected animals within paired genotypes with two-way ANOVA and Bonferroni’s correction. (**D**) Time spent on open arms in elevated plus maze assay at 6 months for WT and PS19 animals injected with AAV9 as in (B). Box and whisker plot represents median, interquartile range, minimum and maximum. n>=9 animals per group. Statistics performed with Welch’s t-test. (**E**) Time spent on closed arms in elevated plus maze assay at 6 months for WT and PS19 animals injected with AAV9 as in (B). Box and whisker plot represents median, interquartile range, minimum and maximum. n>=9 animals per group. Statistics performed with Welch’s t-test. (**F**) Distance moved in elevated plus maze assay at 6 months for WT and PS19 animals injected with AAV9 as in (B). Box and whisker plot represents median, interquartile range, minimum and maximum. n>=9 animals per group. Statistics performed with Welch’s t-test. (**G**) Percent freezing during conditioning period in fear conditioning assay at 6 months for WT and PS19 animals injected with AAV9 as in (B). Data represents mean and SEM. n>=3 animals per group. Statistics performed between GFP and Ser_42_-GFP injected animals within paired genotypes with two-way ANOVA and Bonferroni’s correction. (**H**) Percent freezing during contextual testing following conditioning in fear conditioning assay. Data represents mean and SEM. n>=3 animals per group. Statistics performed with Mann-Whitney test. (**I**) Percent freezing during periods of cued stimulus (CS) following conditioning in fear conditioning assay. Data represents mean and SEM. n>=3 animals per group. Statistics performed with Mann-Whitney test.

A comparison of Ser_42_-GFP injected mice with GFP controls showed polySer induced toxicities in both WT and PS19 animals. Notably, Ser_42_-GFP mice had reduced weight relative to GFP-injected animals of the same genotype (Figure 1B). Furthermore, we observed Ser_42_-GFP injected mice display abnormal motor behaviors such as gait abnormalities, head tilt, and circling phenotypes (Supplemental Movies 1-3). Reflecting these defects in locomotor function, Ser_42_-GFP injected mice had reduced performance on the rotarod assay in both WT and PS19 mice from four to six months of age (Figure 1C).

Additional behavioral assays also highlighted differences in Ser_42_-GFP injected animals. In the elevated plus maze assay, Ser_42_-GFP mice spent increased time in open arms and reduced time in closed arms relative to GFP-injected WT or PS19 animals, while the total distance moved was not significantly different (Figure 1D-F). This suggests Ser_42_-GFP injected mice have reduced natural aversion. In the open field assay, GFP or Ser_42_-GFP expressing mice had no significant differences in the time spent in the center (Figure S1A,B). Still, WT Ser_42_-GFP injected animals showed an increase in the total distance moved relative to controls (Figure S1C). Lastly, we tested whether Ser_42_-GFP expression affected behavior in a fear conditioning assay. We observed Ser_42_-GFP mice show reduced freezing during conditioning (Figure 1G). This reduced freezing is also evident during subsequent testing with either contextual or cued stimuli (Figure 1H,I). However, we also observed reduced freezing during the period prior to cued stimuli suggesting Ser_42_-GFP mice have increased activity at baseline (Figure S1D).

These results collectively show that AAV9-mediated expression of polySer in the CNS leads to motor and behavioral dysfunction in both WT and tau transgenic animals.

### Polyserine overexpression induces Purkinje cell loss

As Ser_42_-GFP expression induced locomotor deficiencies, we next looked for morphological changes that could underlie the observed phenotypes. Of particular interest were Purkinje cells due to their critical role in coordinating movement and the strong tropism of AAV9.^12,13^

We performed immunostaining for GFP and Pcp2 – a Purkinje cell marker – on cerebellum sections from GFP and Ser_42_-GFP injected WT or PS19 transgenic animals. Consistent with previous reports, we observed high transduction of Purkinje cells in GFP-injected controls (Figure 2A).^13^ Strikingly, there was a dramatic loss of Purkinje cells in Ser_42_-GFP injected WT and PS19 mice relative to controls at four months of age (Figure S2A). By six months of age, Purkinje cells were nearly completely lost in Ser_42_-GFP expressing mice (Figure 2A). We measured the cerebellum area and observed a significant reduction in Ser_42_-GFP injected animals relative to controls which became more pronounced over time, consistent with observed loss of Purkinje cells (Figure 2B,C). In contrast, hippocampal neurons – also highly transduced by AAV9 – did not show notable loss in Ser_42_-GFP injected animals (Figure S2B). This suggests Purkinje cells may express higher levels of transgene and/or be particularly susceptible to polySer-induced toxicity.

**Figure 2.**
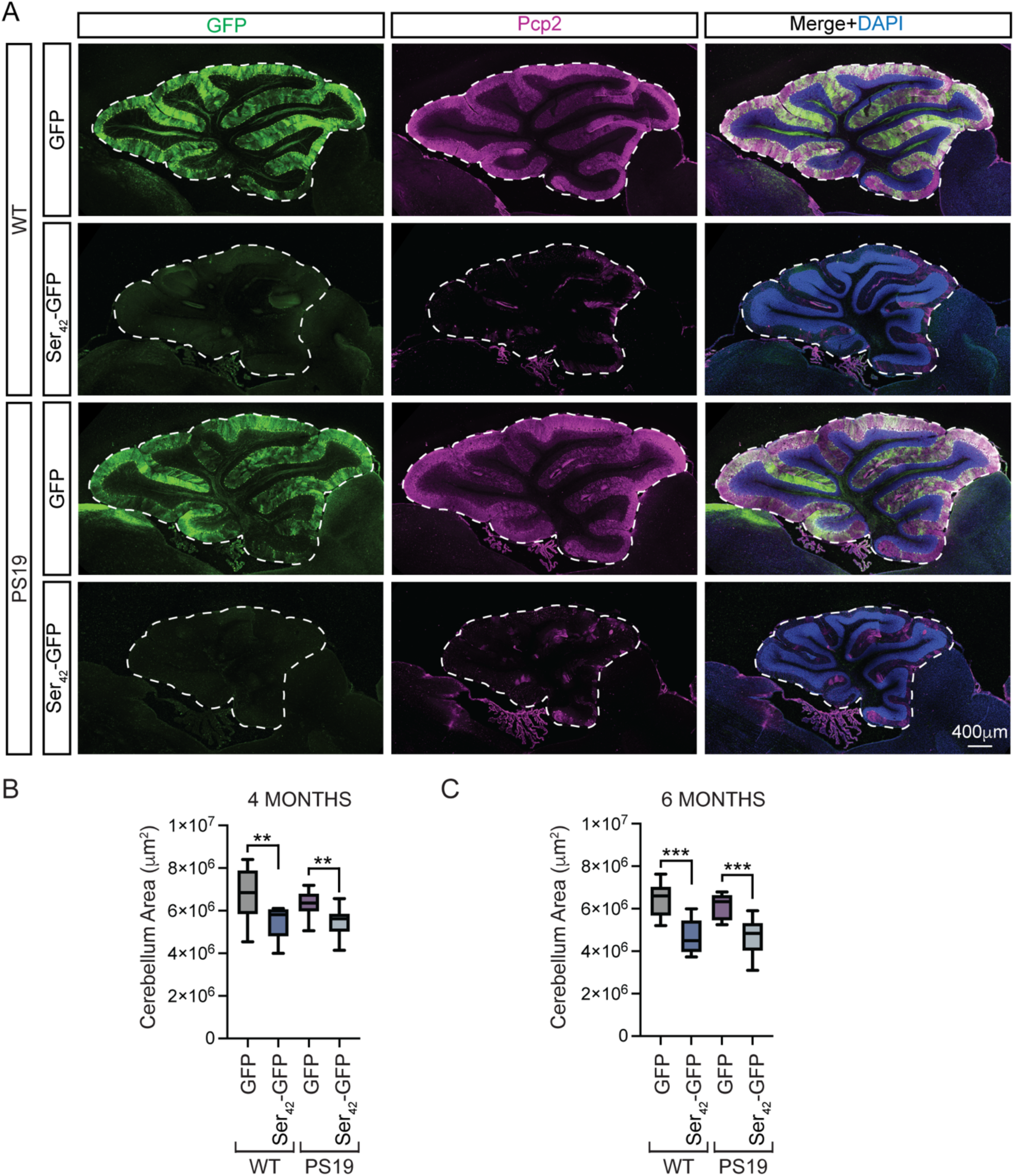
Polyserine expression induces Purkinje cell loss. **(A)** Immunostaining of DAPI (blue), GFP (green) and Pcp2 (magenta) in the cerebellum of WT and PS19 tau transgenic animals at 6 months of age injected with 1×10^11^ vgs of AAV9-GFP or AAV9-Ser_42_-GFP per animal by ICV injection at P1. (**B**) Quantification of cross-sectional area of the cerebellum of WT and PS19 animals at 4 months of age injected with AAV9 as in (A). See Figure S2A. Box and whisker plot represents median, interquartile range, minimum and maximum. n>=8 sections from n>=3 animals per group. Statistics performed with Mann-Whitney test. (**C**) Quantification of cross-sectional area of the cerebellum of WT and PS19 animals at 6 months of age injected with AAV9 as in (A). Box and whisker plot represents median, interquartile range, minimum and maximum. n>=9 sections from n>=3 animals per group. Statistics performed with Mann-Whitney test.

Thus, polySer overexpression causes Purkinje cell loss, likely underlying some of the observed locomotor abnormalities.

### Polyserine forms assemblies in mouse neurons

We previously observed that exogenous polySer assemblies in cultured cells are preferred sites of tau aggregation.^7^ To determine the localization of Ser_42_-GFP in neurons *in vivo*, we performed immunostaining of brain sections from AAV9-injected mice at two months of age. Consistent with our previous findings in cell models, we observed that Ser_42_-GFP formed assemblies within CA1 and DG hippocampal neurons while GFP alone maintained diffuse localization throughout transduced cells (Figure 3A,B). A similar localization was observed in Purkinje cells when examined before their significant loss (Figure 3C). Analysis of PS19 animals also exhibited punctate assemblies of polySer (Figure S3A-C). Thus, these findings, along with reports from other groups of polySer localization in SCA8 and HD, underline the self-assembly properties of polySer.^8,9^

**Figure 3.**
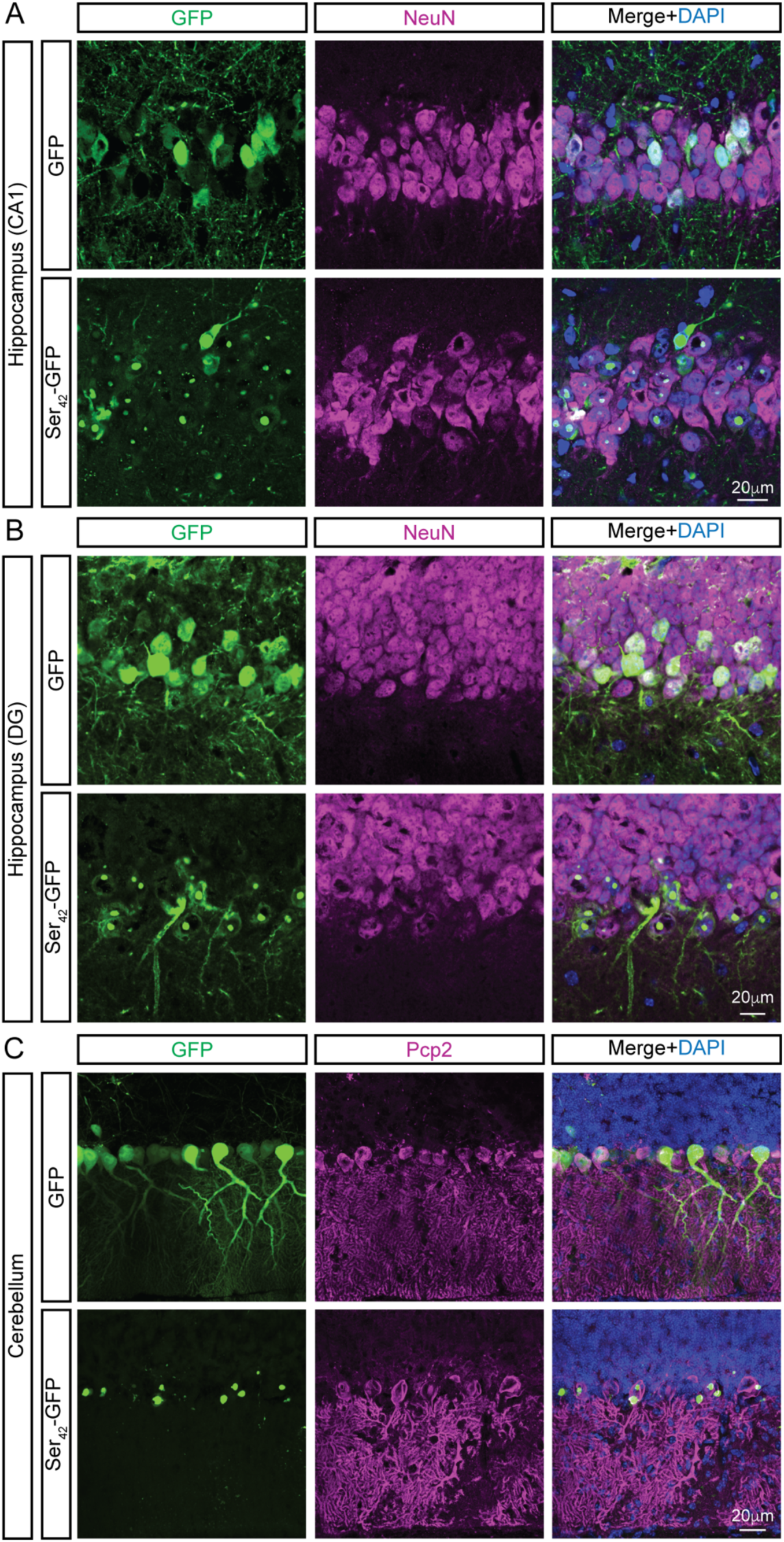
Polyserine forms assemblies in neurons *in vivo*. **(A)** Immunostaining of DAPI (blue), GFP (green) and NeuN (magenta) in CA1 hippocampal neurons of WT animals at 2 months injected with 2×10^11^ vgs of AAV9-GFP or AAV9-Ser_42_-GFP per animal. (**B**) Immunostaining of DAPI (blue), GFP (green) and NeuN (magenta) in dentate gyrus (DG) hippocampal neurons of WT animals at 2 months injected with AAV9 as in (A). (**C**) Immunostaining of DAPI (blue), GFP (green) and Pcp2 (magenta) in Purkinje cells of WT animals at 2 months injected with AAV9 as in (A).

### Polyserine exacerbates tau pathology in PS19 mice

We next examined whether Ser_42_-GFP altered the progression of tau pathology in PS19 mice. Notably, delivery of 1×10^11^ viral genomes (vgs) (1X) of Ser_42_-GFP resulted in early fatalities in PS19 mice but not in WT controls (Figure 4A). Further, delivery of a higher dose of 2×10^11^ vgs (2X) of Ser_42_-GFP resulted in a significant reduction in survival of PS19 mice relative to WT animals (Figure 4A). These findings demonstrate that the expression of Ser_42_-GFP affects lifespan more severely in mice expressing a pathogenic human tau transgene.

**Figure 4.**
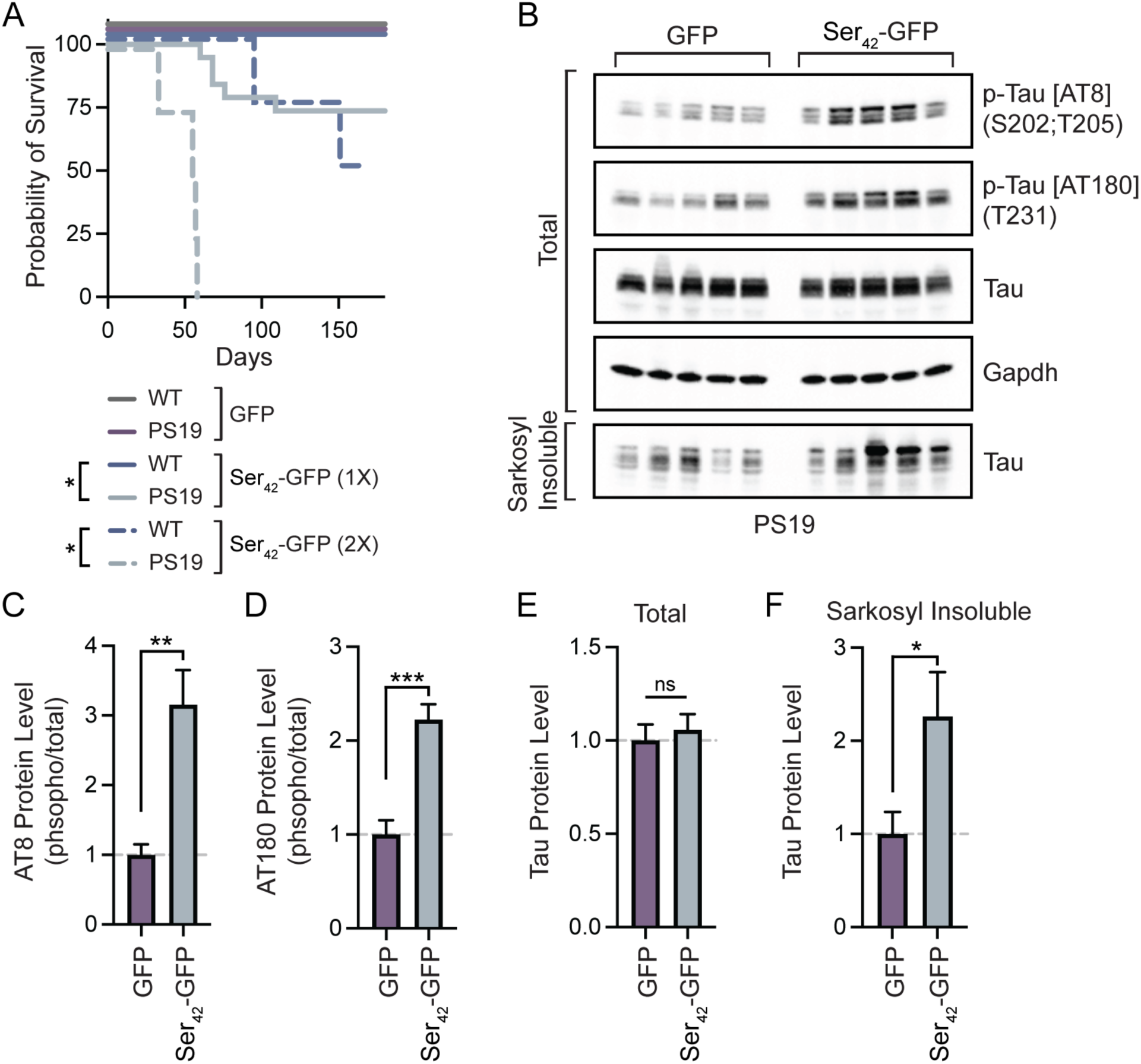
AAV9-mediated overexpression of polyserine exacerbates tau pathology in transgenic mice. (**A**) Kaplan-Meier plot of the survival of WT and PS19 tau transgenic animals injected with GFP or Ser_42_-GFP at 1×10^11^ vgs (1X) (n>=12 animals per group) or 2×10^11^ vgs (2X) (n>=4 animals per group) per animal at P1. Statistics performed with Mantel-Cox test. (**B**) Western blot of phosphorylated tau (p-tau) (AT8), p-tau (AT180), total tau and GAPDH in total or sarkosyl insoluble extracts from PS19 animals injected with 1×10^11^ vgs of AAV9-GFP or AAV9-Ser_42_-GFP at P1. (**C**) Quantification of AT8 p-tau protein levels normalized to total tau as in (B). Data represents mean and SEM. n=5 animals per group. Statistics performed with unpaired t-test. (**D**) Quantification of AT180 p-tau protein levels normalized to total tau as in (B). Data represents mean and SEM. n=5 animals per group. Statistics performed with unpaired t-test. (**E**) Quantification of total tau protein levels normalized to GAPDH as in (B). Data represents mean and SEM. n=5 animals per group. Statistics performed with unpaired t-test. (**F**) Quantification of total tau levels in sarkosyl insoluble fractions normalized to GFP-injected control as (B). Data represents mean and SEM. n=5 animals per group. Statistics performed with unpaired t-test.

We next monitored direct readouts of tau pathology by Western blot. We observed that brain extracts from Ser_42_-GFP injected PS19 animals had increased levels of phosphorylated tau (p-tau) (monitored by AT8 and AT180 antibodies) relative to GFP controls (Figure 4B-D). Furthermore, while there were no significant changes in total tau levels across groups, Ser_42_-GFP expressing mice had increased sarkosyl insoluble tau (Figure 4B, E, F). Next, we performed immunostaining to measure p-tau levels specifically in the hippocampus with an antibody for Ser202/Thr205 p-tau (AH36). As expected, we observed an increase in the p-tau positive area of the hippocampus in GFP-injected PS19 mice compared to WT (Figure 5A,B). Further, consistent with Western blotting results, we also observed a significant increase in p-tau staining in Ser_42_-GFP PS19 mice as compared to GFP (Figure 5A,B).

**Figure 5.**
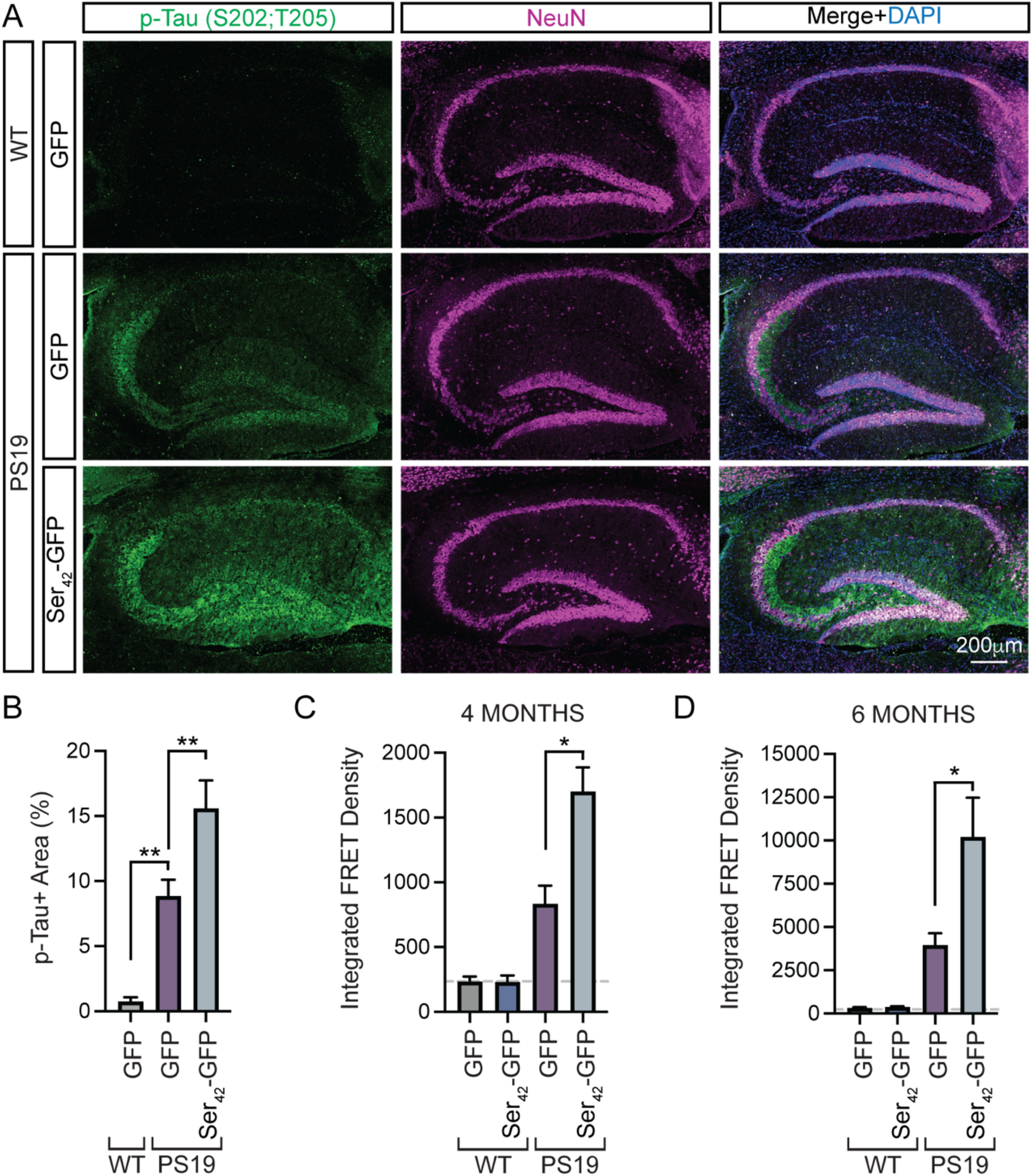
AAV9-mediated overexpression of polyserine increases tau seeding capacity. (**A**) Immunostaining of DAPI (blue), p-Tau (AH36) (green) and NeuN (magenta) in the hippocampus of WT or PS19 at 6 months injected with 1×10^11^ vgs of AAV9-GFP or AAV9-Ser_42_-GFP. (**B**) Quantification of the percent area of the hippocampus positive for AT8 staining as in (A). Data represents mean and SEM. n>=11 sections from n>=4 animals per group. Statistics performed with one-way ANOVA. (**C**) Seeding capacity of sarkosyl insoluble fractions from brain extracts of WT and PS19 mice at 4 months of age injected with GFP or Ser_42_-GFP. Data represents mean and SEM. n=3 animals per group. Statistics performed with Welch’s t-test. (**D**) Seeding capacity of sarkosyl insoluble fractions from brain extracts of WT and PS19 mice at 6 months of age injected with GFP or Ser_42_-GFP. Data represents mean and SEM. n>=4 animals per group. Statistics performed with Welch’s t-test.

We also assessed the burden of tau pathology by measuring seeding activity. To do so, we transfected total and sarkosyl fractionated brain extracts from AAV9-injected animals into HEK293 tau biosensor cells that express the tau repeat domains with a P301S mutation fused to a FRET pair (CFP/YFP), which enables quantification of tau aggregation as a measure of seeding capacity by flow cytometry.^14^ Importantly, we observed a baseline level of FRET signal in sarkosyl soluble fractions from WT or PS19 mice, validating our fractionation and assay (Figure S4A,B). Strikingly, although levels of aggregation were low at four months of age, there was a significant increase in the seeding capacity of sarkosyl insoluble fractions of brain extracts from Ser_42_-GFP tau transgenic mice relative to GFP controls (Figure 5C). Further, at a later stage of six months of age, we once again observed a significant increase in the seeding capacity of sarkosyl insoluble extracts from Ser_42_-GFP expressing tau transgenic animals (Figure 5D).

Collectively, these results demonstrate that Ser_42_-GFP can exacerbate measures of tau pathology in a transgenic mouse model.

## Discussion

This study provides several observations that polySer can promote tau aggregation and pathology in the mammalian brain. First, polySer expression significantly reduces lifespan in tau transgenic mice (Figure 4). Second, we observe that polySer increases levels of phosphorylated and insoluble tau in the brain (Figures 4 & 5). Finally, we observe that polySer brains have greater levels of seeding competent tau (Figure 5). Together, these observations demonstrate polySer levels can impact tau pathology in mice, consistent with earlier work showing the levels of polySer expression correlate with the extent of tau aggregation in cell lines.^7^

In principle, increases in polySer levels could increase tau aggregation by multiple mechanisms. Since we observe polySer forms cytoplasmic assemblies that are a preferred site for tau aggregation in cellular models, polySer could similarly increase tau fibrillization *in vivo.*^7^ In support of this model, we observe that viral-mediated expression of polySer in neurons also forms assemblies (Figure 3). Other groups have observed that polySer forms aggregates *in vitro*, puncta in C. elegans and assemblies in the brains of SCA8 and HD patients.^8,9,15–17^ Nonetheless, we cannot exclude that polySer may increase tau pathology through indirect mechanisms such as increasing neuroinflammation and/or altering protein homeostasis pathways.

The observation that increases in polySer exacerbates tau markers in a mouse model supports the hypothesis that mislocalization of nuclear polySer domain containing RNA binding proteins to the cytoplasm can promote tau pathology in human disease (Figure 6). This hypothesis is based on the finding that SRRM2 – which contains a polySer domain – mislocalizes to cytosolic tau aggregates in tauopathy cell models, animal models and human post-mortem tissues.^6,7^ Moreover, while typically present in nuclear speckles, SRRM2 and the polySer-rich protein PNN define cytoplasmic assemblies, which are sites of tau aggregation and are increased in iPSC derived neurons under inflammatory conditions.^7^ Further, as reduced nuclear import of SRRM2 was identified in a mouse model with Aβ pathology, one possibility is that increased cytoplasmic mislocalization of SRRM2 – and potentially other polySer domain-containing proteins – may sufficiently increase the cytoplasmic pool of polySer to accelerate the development of tau pathology and contribute to disease.^18^

**Figure 6.**
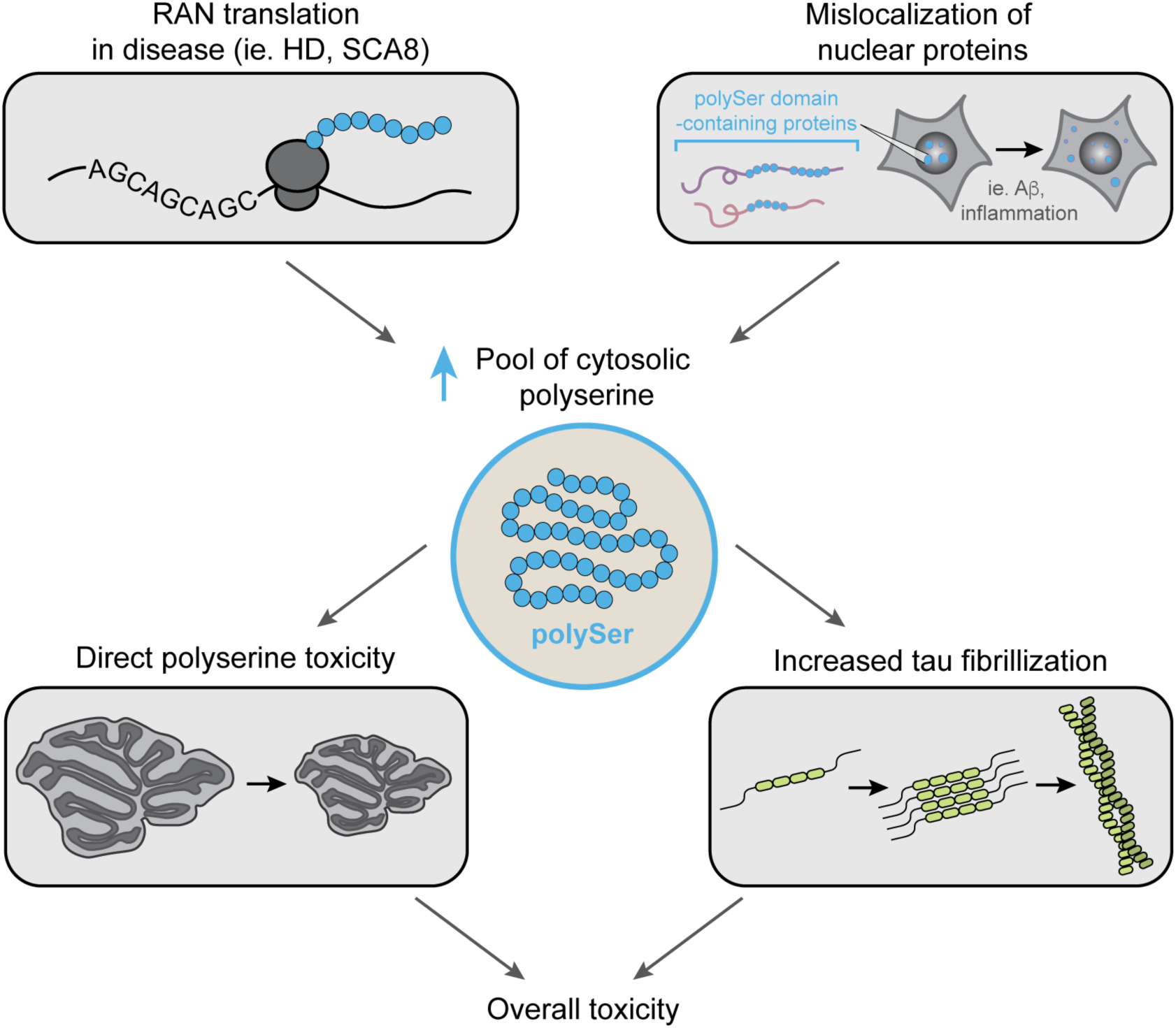
Model of polyserine sources and potential contributions to toxicity. PolySer can be expressed due to RAN translation in repeat expansion diseases such as SCA8 and HD. Cytosolic levels of polySer can also increase due to polySer-domain containing nuclear proteins mislocalizing in response to stressors such as Aβ or inflammation. Increases in the pool of polySer can have direct deleterious effects such as the loss of Purkinje cells which occurs through unknown mechanisms and/or promote tau fibrillization and pathology. The additive effect of such pathways amounts to the overall toxicity induced by polySer.

Independent of effects on tau pathology, we also demonstrate that excess polySer is toxic to neurons. A key observation is that AAV9-mediated expression of polySer causes locomotor defects and aberrant behavior in WT mice (Figure 1). Moreover, this is coincident with a loss of Purkinje cells, which likely contributes to behavioral defects (Figure 2). Interestingly, a previous study pointed to toxicity of exogenously delivered polySer peptides leading to reduced viability of cells *in vitro*, vacuolar degeneration, and behavioral changes following direct delivery to the ventricles of mice.^15^ Collectively, these findings underline a general toxic role for polySer *in vivo*.

Importantly, polySer expression has been identified in multiple disease contexts where it may contribute to pathobiology. Most notably, polySer is detectable in patient tissues from SCA8 and HD cases which are caused by CTG and CAG repeat expansions, respectively.^8,9^ Due to bidirectional transcription and RAN translation, polySer can be expressed by translation in the AGC frame. As a result, polySer could be expressed in additional repeat expansion disorders caused by CTG/CAG expansions such as SCA types 1, 2, 3, 6, 7, 12 and 17, spinal and bulbar muscular atrophy (SBMA) and others. Consistent with this, polySer RAN translation products are present in SCA type 12 iPSC-derived cell lines.^19^ Accumulations of polySer in SCA8 post-mortem tissues are mainly detected in white matter regions where they correlate with demyelination and axonal degeneration.^8^ While our viral-delivered polySer is expressed primarily in neurons, it is sufficient to induce locomotor dysfunction in mice underlaid by a dramatic loss of Purkinje cells which are the primary cell type affected in spinocerebellar ataxias. Taken together, these observations support a potential contribution of polySer toxicity to SCA8 disease progression and potentially other diseases.

Our results raise the possibility that polySer could also contribute to SCA8 and HD disease progression in a tau-dependent manner. Specifically, as polySer can exacerbate tau aggregation, it may do so when expressed in the context of SCA8 or HD.^8,9^ Consistent with this hypothesis, HD patients develop tau pathology, although it is not yet clear how it may impact disease progression.^20^ Moreover, a small study of post-mortem tissues from SCA8 patients revealed two of four patients studied had extensive tauopathy, while another SCA8 patient presented with biomarkers of PSP, a well-characterized tauopathy.^21,22^ These observations reveal the intriguing possibility that patients with a repeat expansion disease that produces polySer may be more likely to develop tau pathology. However, addressing this issue will require substantial examination of patients for both polySer expression and tau pathology. Thus, developing our understanding of the links between polySer and tau has the potential to provide insight into disease mechanisms beyond primary tauopathies.

## Acknowledgements

M.V.A. is a Howard Hughes Medical Institute Awardee of the Life Sciences Research Foundation. This work was supported by funds to R.P. from the Howard Hughes Medical Institute. C.A.H is supported by funding from NIH AG064465, AG083268, and DA055781. This study utilized resources provided by the University of Colorado Boulder Department of Biochemistry Cell Culture Facility (RRID SCR_19309) and Flow Cytometry Core (RRID SCR_018988).

## Author Contributions

M.V.A. and R.P designed the study. M.V.A. performed experiments. V.L.N performed mouse behavioral studies. C.A.H. supervised mouse studies. M.V.A. and R.P wrote and revised manuscript with input from all authors.

## Methods

### Animal procedures

All work with mice was performed in accordance with the National Institutes of Health Guide on the Care and Use of Animals and approved by the Institutional Animal Care and Use Committee of University of Colorado Boulder. PS19 tau transgenic mice on a C57BL/6-congenic background were obtained from Jackson mice (Strain #024841)^11^ and crossed with C57BL/6J (Strain #000664) to obtain wild-type or heterozygous experimental animals. Genotyping was performed using DNA extracted from tails with primers listed in Table 2. Aggregated data of male and female animals is presented.

**Table 1.** Antibodies.

| Target | Manufacturer | Catalog # |
| --- | --- | --- |
| DAPI | Invitrogen | 62248 |
| Mouse anti-Pcp2 | Sigma | sc-137064 |
| Chicken anti-GFP | Aves | GFP-1020 |
| Guinea pig anti-NeuN | Synaptic Systems | 266 004 |
| Rabbit anti-tau (K18) | Abcam | Ab218314 |
| Mouse anti p-tau (AT8) | Thermo Scientific | MN1020 |
| Mouse anti p-tau (AT180) | Thermo Scientific | MN1040 |
| Mouse anti-GAPDH | Millipore Sigma | MAB374 |
| Rabbit anti p-tau (AH36) | Stress Marq | SMC-601D |
| Anti-rabbit HRP | Cell Signaling | 7074S |
| Anti-mouse HRP | Cell Signaling | 7076S |
| 488 anti-chicken | Jackson Labs | 703-545-155 |
| CY3 anti-rabbit | Jackson Labs | 711-165-152 |
| CY3 anti-mouse | Jackson Labs | 715-165-150 |
| CY5 anti-guinea pig | Jackson Labs | 706-175-148 |

**Table 2.**
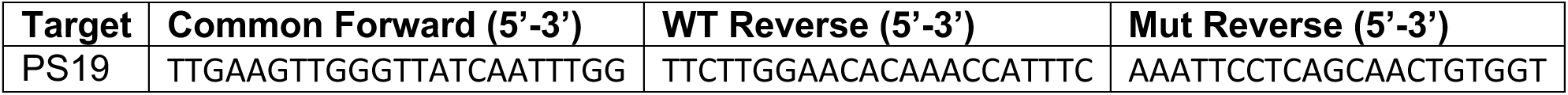
Genotyping Primers.

### AAV9 production and delivery

The ORFs of GFP and Ser_42_-GFP were cloned into the pAAV-CMV plasmid (Addgene #105530). Endofree plasmids were prepped with ZymoPURE Midiprep plasmid kit (Zymo Research) and AAV9 were generated by Vector BioLabs. AAV9 were delivered by a single injection to the right lateral ventricle at a dose of 1×10^11^ or 2×10^11^ genome copies per animal at P1 in a PBS solution containing FastGreen dye (Sigma) as previously described.^23^

### Mouse behavioral assays

Mice were tested on the accelerating rotarod assay as previously described.^24^ Briefly, animals were trained one week prior to first testing at 4 months of age and subsequent testing was performed monthly. A 60 second warm up was performed before 3 trials which consisted of 5 minutes of acceleration from 4 to 40 rotations per minute. The average of 3 trials was plotted for each animal.

For elevated plus maze assay, animals were moved to testing room and singly housed with 55dB white noise prior to testing. After 1 hour of acclimation, mice were placed on the elevated plus maze for 5 minutes and monitored with EthoVision XT video tracking software. The percentage of time spent on either open or closed arms relative to the total time of testing in assay was reported.

For open-field assay, animals were moved to testing room and singly housed with 55dB white noise prior to testing. Following 1 hour of acclimation, animals were placed in the arena for a total of 10 minutes and monitored with EthoVision XT video tracking software. The percentage of time spent in each portion of the arena relative to the total testing time was reported.

Fear conditioning assay was performed as previously described.^25,26^ Briefly, animals were acclimated for 1 hour prior to testing singly housed with 55dB white noise. Animals were monitored in isolation cubicles (30” W x 17.75” D x 18.5”H) (Coulbourn) with FreezeFrame software (Actimetrics) during all testing. During training, animals were subject to two pairings of a tone (30 seconds, 85-dB white noise) and foot-shock (2 seconds, 0.5mA). Tone was (120-150 seconds and 210-240 seconds) followed by foot-shock (148-150 seconds and 238-240 seconds) two times during a total session length of 5 minutes. For the training period the house lights were on and peppermint odor present. The following day, mice were again acclimated prior to memory tests. Cued and contextual testing were performed in the same day with a minimum of 1 hour between testing for each animal. The order of cued or contextual testing was randomized. During context testing, animals were returned to the same isolation cubicles and exposed to the same conditions as the training for a total of 5 minutes. During cued testing, animals were placed in randomized isolation cubicles with red light on, infrared light on, house lights off, white acrylic floor over shock grid, inserts with different display patterns over test cage walls and vanilla odor. Animals were monitored with these conditions for a total of five minutes with tone at the same times as in the training test but no paired foot shock. The average percent of time freezing during each 10 second interval of training was reported. For context testing, the average time spent freezing across the 5 minute test was reported. For the baseline freezing measured prior to the cued stimulus, the average time spent freezing in cued testing prior to playing of the first tone (0-120 seconds) was reported. For cued testing, the average time spent freezing during both periods of tone playing was reported.

### Immunohistochemistry

Following dissection brains were hemisected and post-fixed in 4% PFA in PBS for 24 hours. Tissue was cryoprotected in 30% sucrose in PBS for 24 hours before flash freezing in O.C.T. sagittal sections were prepared on the cryostat at 30um thickness and mounted on charged slides. Staining was performed in a humidified chamber. Sections were blocked with 10% donkey serum in PBS with 0.4% Triton-X at room temperature then incubated with primary antibodies overnight at 4°C in blocking buffer. Following primary, slides were washed 3 x 10 minutes in PBS with 0.4% Triton-X then incubated with secondary antibodies for 3 hours at room temperature in blocking buffer. Following secondary, slides were washed 3 x 10 minutes in PBS before mounting with Fluoromount G.

### Confocal Imaging and Image Quantification

Imaging of immunostaining was performed on a Nikon spinning disk confocal microscope with a 10X (NA 0.45) objective and 3μm steps using stitching and overlap to obtain composite images of the entire cerebellum or hippocampal cross-section. Images of polySer assemblies were acquired with a 40X (NA 1.15) objective. Quantification of the cross-sectional area of cerebellum was performed with manual ROI selection and NIS-Elements AR (Nikon) software. Quantification of p-tau positive area of the hippocampus was performed with a CellProfiler pipeline. Specifically, maximum projections of scans were manually annotated and masked for the hippocampus based on NeuN staining, then p-tau positivity was defined by intensity thresholding. The percent positive area was determined and calculated relative to the total hippocampal area of each section.

### Sarkosyl Fractionation

Sarkosyl fractionation was performed as previously reported by others.^27^ Briefly, flash frozen brain tissue was homogenized with a dounce homogenizer in 10mM Tris-HCl (pH 7.4), 0.8M NaCl and 1mM EDTA buffer supplemented with protease and phosphatase inhibitors (Roche) at a weight to volume ratio of 1:10. Lysate was centrifuged at 21,000g for 10 minutes at 4°C. The supernatant was incubated at room temperature with 1% sarkosyl for 1 hour then centrifuged for 1 hour at 186,000g at 4°C. Pellet was washed once with PBS then centrifuged again for 1 hour at 186,000g at 4°C. Pellet was resuspended in PBS at 1/2 the starting total volume of lysate.

### Western blotting

Western blotting was performed as previously described.^7^ Samples were boiled in 2X Laemmli sample buffer at 95°C for 10 minutes then run on pre-cast 4-20% Tris-glycine gels (Invitrogen). Samples were transferred to nitrocellulose membrane using iBlot 2 gel transfer device (Invitrogen). Samples were blocked with 5% milk or 5% BSA (for phosphor blots) in TBS-T (0.1%Tween) for 1 hour, incubated with primary antibodies for 2 hours, washed 3 x 10 minutes with TBS-T, incubated with secondary antibodies in TBS-T for 1 hour, then wased 6 x 5 minutes with TBS-T. All steps were performed at room temperature. Blots were developed with Clarity Western ECL Substrate (Bio-rad) and imaged with iBright imaging system (Thermo Fisher).

### Cell Culture

HEK293T tau biosensor cells were purchased from ATCC and culture in DMEM with 10% FBS, 0.2% penicillin-streptomycin at 37°C with 5% carbon dioxide.^14^ Brain homogenate was transfected into cells with Lipofectamine3000. For total and sarkosyl soluble extracts 1μl of sample was transfected per 24-well 24 hours after plating at a density of 100K cells/well. For sarkosyl insoluble extracts 20μl of sample was transfected per 24-well 24 hours after plating at a density of 100K cells/well.

### Flow Cytometry

Flow cytometry was performed as previously reported.^7^ In brief, cells were trypsinized, washed with PBS and passed through 40μm sterile filters then flow cytometry was done with a BD FACSCelesta^TM^ Cell Analyzer using filter sets 405-450 (CF) and 405-525 (FRET). Analysis was performed for FlowJo and control samples were set to a false FRET percentage of 1 as previously reported.^7^ Reported integrated FRET density was calculated as the produce of the percentage of FRET-positive cells and median fluorescence intensity.

### Statistical Analysis

Statistical tests and replicates are reported in figure legends. In figures, statistical significance is reported as follows (*) p<0.05, (**) p<0.01, (***) p<0.001, (****) p<0.0001 for the defined p-values.

---

**Figure S1.**
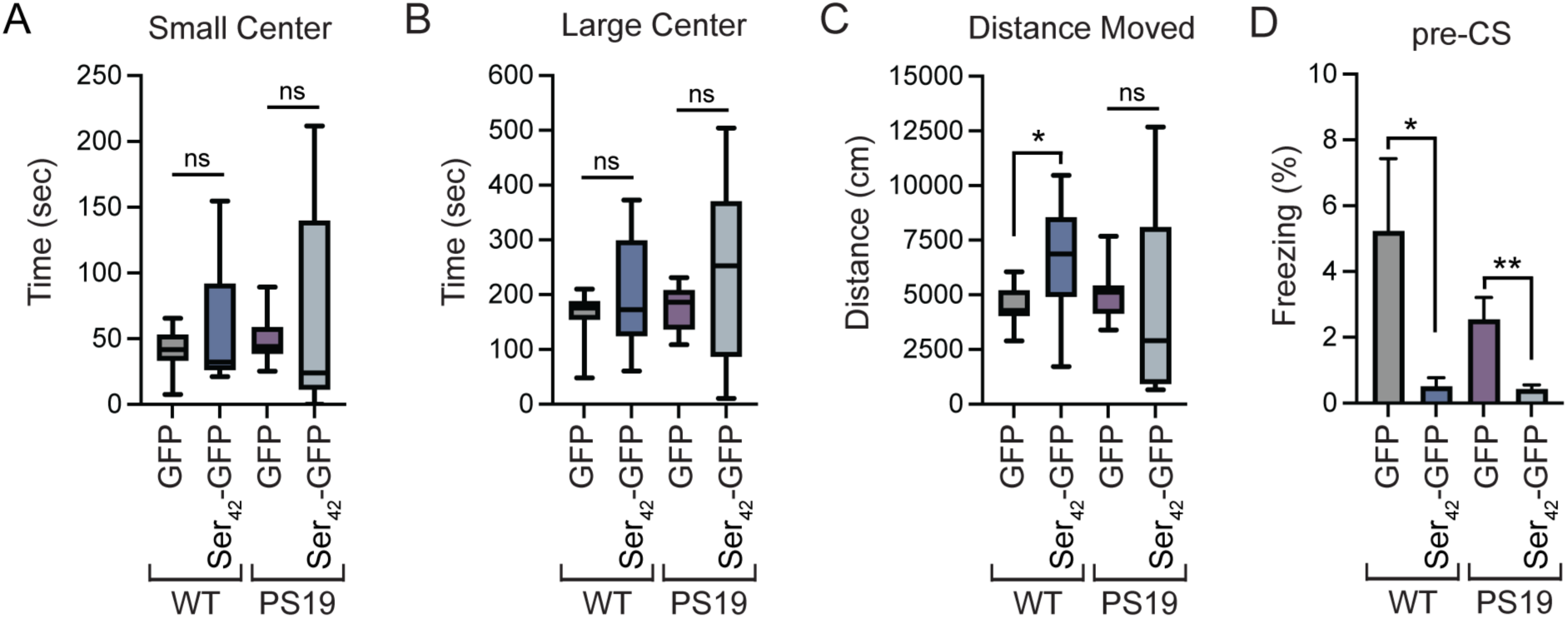
AAV9-mediated overexpression of polyserine leads to motor dysfunction in wild-type and tau transgenic mice. (**A**) Time spent in the small center during open-field assay at 6 months for WT and PS19 tau transgenic animals injected with AAV9-GFP or AAV9-Ser_42_-GFP. Box and whisker plot represents median, interquartile range, minimum and maximum. n>=9 animals per group. Statistics performed with Welch’s t-test. (**B**) Time spent in the large center during open-field assay at 6 months for WT and PS19 tau transgenic animals injected with AAV9 as in (A). Box and whisker plot represents median, interquartile range, minimum and maximum. n>=9 animals per group. Statistics performed with Welch’s t-test. (**C**) Distance moved in open-field assay at 6 months for WT and PS19 animals injected with AAV9 as in (A). Box and whisker plot represents median, interquartile range, minimum and maximum. n>=9 animals per group. Statistics performed with Welch’s t-test. (**D**) Percent freezing during period prior to cued stimulus (pre-CS) following conditioning in fear conditioning assay. Data represents mean and SEM. n>=3 animals per group. Statistics performed with Mann-Whitney test.

**Figure S2.**
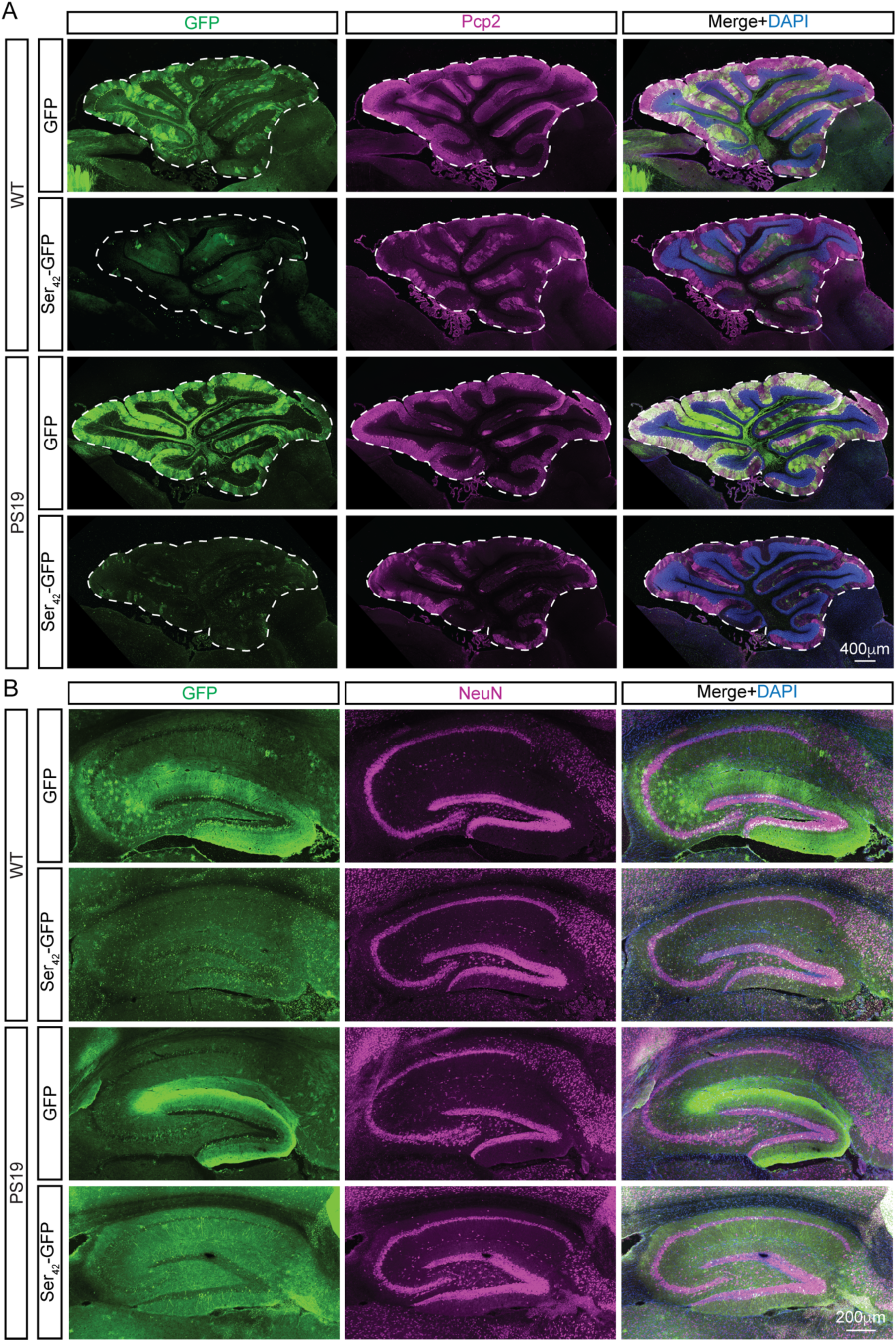
Polyserine expression induces Purkinje cell loss. (**A**) Immunostaining of DAPI (blue), GFP (green) and Pcp2 (magenta) in the cerebellum of WT and PS19 animals at 4 months of age injected with 1×10^11^ vgs of AAV9-GFP or AAV9-Ser_42_-GFP at P1 by ICV injection. (**B**) Immunostaining of DAPI (blue), GFP (green) and NeuN (magenta) in the hippocampus of WT and PS19 animals at 6 months of age injected with 1×10^11^ vgs of AAV9-GFP or AAV9-Ser_42_-GFP at P1 by ICV injection.

**Figure S3.**
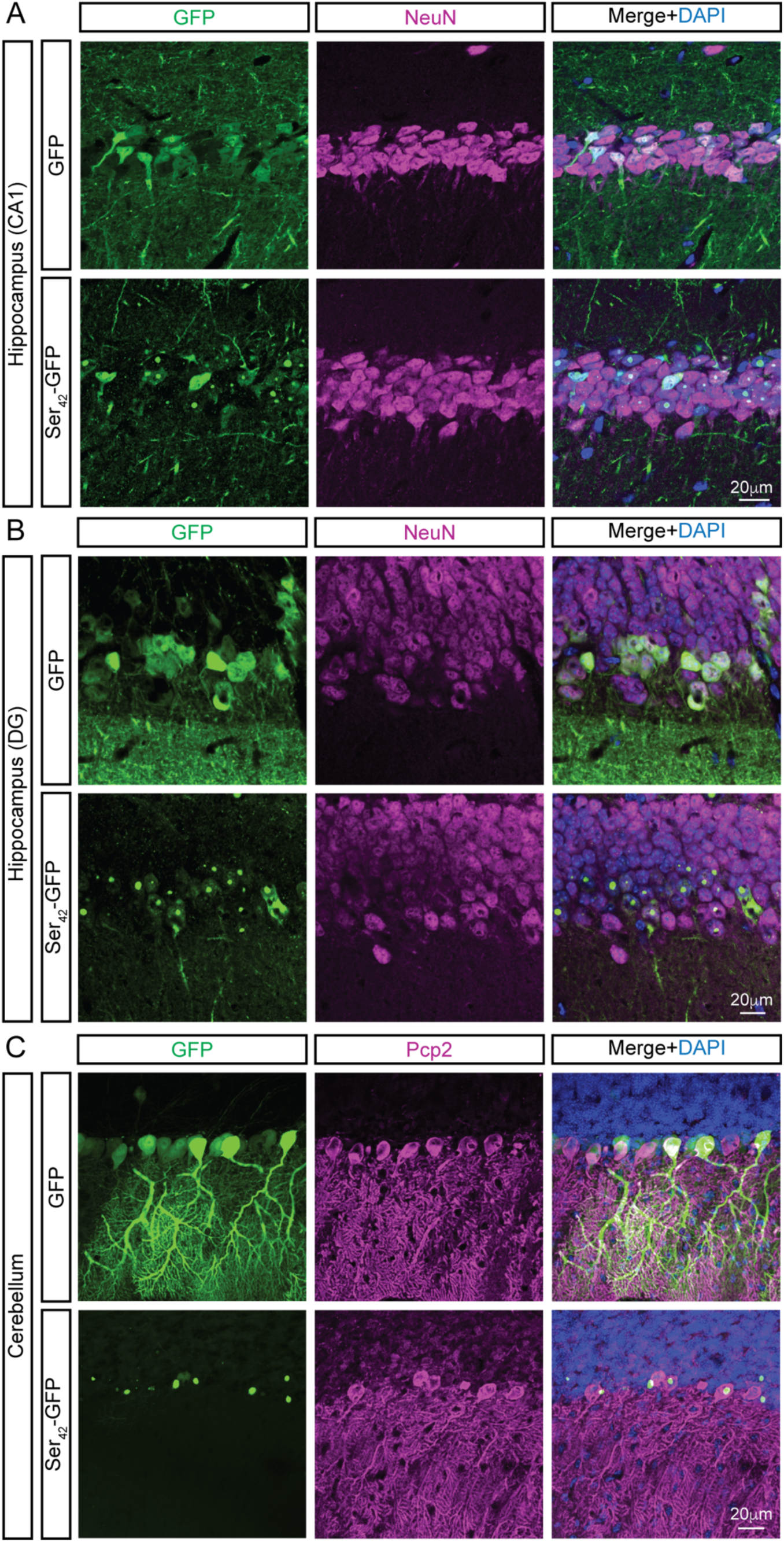
Polyserine forms assemblies in neurons of tau transgenic mice. (**A**) Immunostaining of DAPI (blue), GFP (green) and NeuN (magenta) in CA1 hippocampal neurons of PS19 animals at 2 months injected with 2×10^11^ vgs of AAV9-GFP or AAV9-Ser_42_-GFP per animal. (**B**) Immunostaining of DAPI (blue), GFP (green) and NeuN (magenta) in DG hippocampal neurons of PS19 animals at 2 months injected with 2×10^11^ vgs of AAV9-GFP or AAV9-Ser_42_-GFP per animal. (**C**) Immunostaining of DAPI (blue), GFP (green) and Pcp2 (magenta) in Purkinje cells of PS19 animals at 2 months injected with 2×10^11^ vgs of AAV9-GFP or AAV9-Ser_42_-GFP per animal.

**Figure S4.**
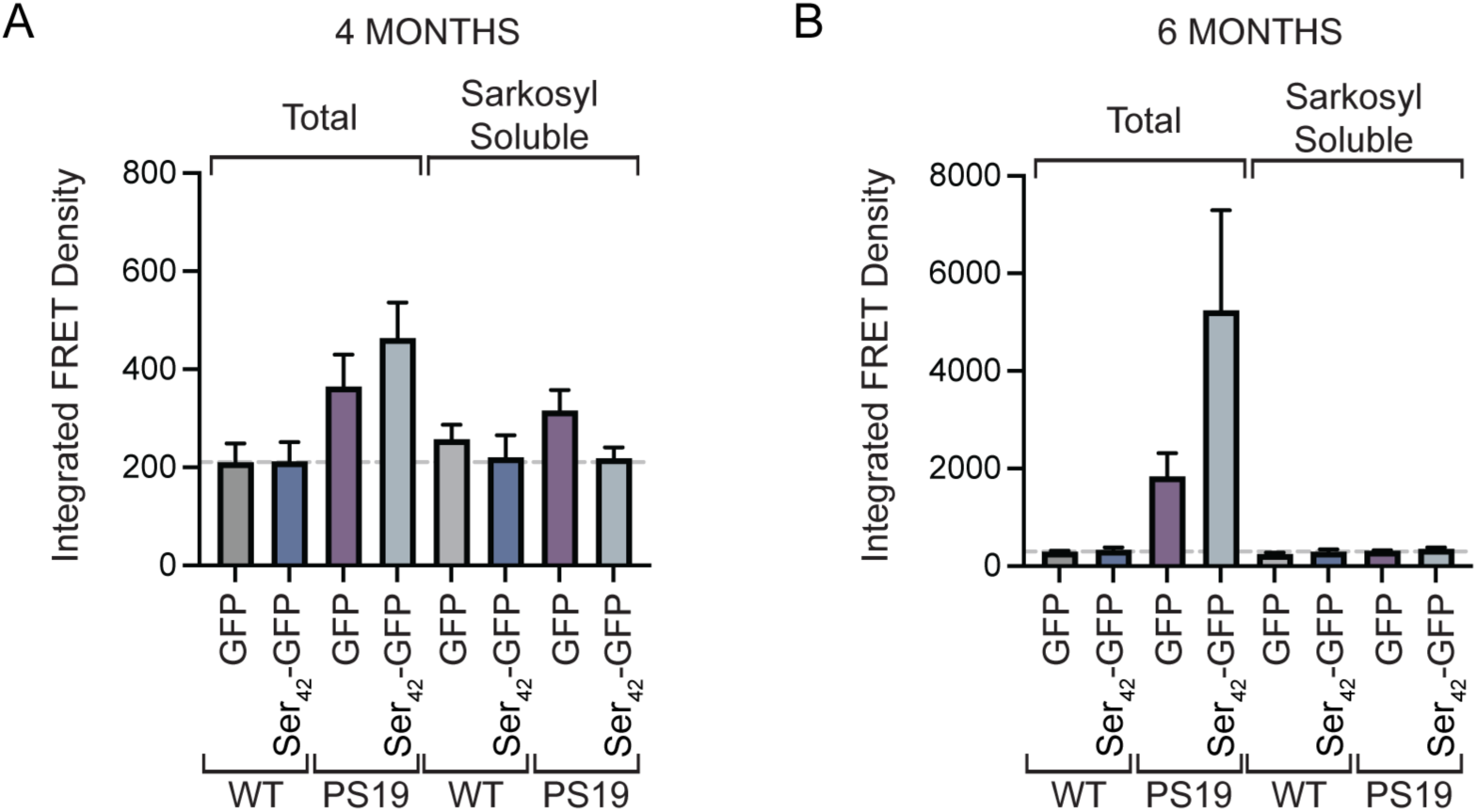
Seeding capacity of total and sarkosyl soluble fractions. (**A**) Seeding capacity of total and sarkosyl soluble fractions from brain extracts of WT and PS19 mice at 4 months of age injected with 1×10^11^ vgs of AAV9-GFP or AAV9-Ser_42_-GFP. Data represents mean and SEM. n=3 animals per group. (**B**) Seeding capacity of total and sarkosyl soluble fractions from brain extracts of WT and PS19 mice at 6 months of age injected with GFP or Ser_42_-GFP as in (A). Data represents mean and SEM. n>=4 animals per group.

**Video 1. Gait abnormality in polySer treated animal.** Gait abnormalities of an AAV9-Ser_42_ treated animal in the right side of the cage at the start of video can be observed relative to a control animal at 6 months of age.

**Video 2. Head tilt in polySer treated animal.** Head tilt phenotype in AAV9-Ser_42_ treated animal at 6 months of age.

**Video 3. Circling phenotype in polySer treated animal.** Circling phenotype in AAV9-Ser_42_ treated animal at 6 months of age.

## Notes

### Competing Interest Statement

The authors have declared no competing interest.

